# pyABC: distributed, likelihood-free inference

**DOI:** 10.1101/162552

**Authors:** Emmanuel Klinger, Dennis Rickert, Jan Hasenauer

## Abstract

Likelihood-free methods are often required for inference in systems biology. While Approximate Bayesian Computation (ABC) provides a theoretical solution, its practical application has often been challenging due to its high computational demands. To scale likelihood-free inference to computationally demanding stochastic models we developed pyABC: a distributed and scalable ABC-Sequential Monte Carlo (ABC-SMC) framework. It implements computation-minimizing and scalable, runtime-minimizing parallelization strategies for multi-core and distributed environments scaling to thousands of cores. The framework is accessible to non-expert users and also enables advanced users to experiment with and to custom implement many options of ABC-SMC schemes, such as acceptance threshold schedules, transition kernels and distance functions without alteration of pyABC’s source code. pyABC includes a web interface to visualize ongoing and finished ABC-SMC runs and exposes an API for data querying and post-processing.

**Availability and Implementation:** pyABC is written in Python 3 and is released under the GPLv3 license. The source code is hosted on https://github.com/neuralyzer/pyabc and the documentation on http://pyabc.readthedocs.io. It can be installed from the Python Package Index (PyPI).

## 1 Introduction

The development of predictive models of biological processes requires the estimation of parameters from experimental data. For model classes such as ordinary differential equations, tailored approaches exploiting specific model properties have been developed to solve these inverse problems (Raue et al. 2015). However, this has not been possible for many other relevant model classes such as stochastic models of intracellular processes or multi-scale models of biological tissues. These models are often so involved and problem-specific that they have to be considered as black-boxes (Jagiella et al. 2017). Black-box models can be simulated but their internal structure cannot be exploited. To parameterize these models, likelihood-free methods, such as ABC-SMC schemes (Sisson et al. 2007; Toni et al. 2009), have been developed. ABC-SMC schemes use numerical simulations of the model to infer its parameters. Although some ABC-SMC frameworks for Python exist, these frameworks either lack customization options like (adaptive) acceptance threshold schedules or transition kernels, (Kangasrääsiö et al. 2016), do not implement scalable parallelization strategies (Jennings and Madigan 2017), only parallelize across the cores of a single machine (Ishida et al. 2015), or only leverage GPUs but not distributed infrastructure (Liepe et al. 2010). To address these shortcomings we developed pyABC.

## 2 Features

### 2.1 Usage

Parameter estimation and model selection for simulator-based, black-box models using several different parallelization strategies is implemented in the pyABC framework. The framework’s features include

- multi-core and distributed execution,
- computation-minimizing and scalable, runtime-minimizing parallelization,
- adaptive, local transition kernels and acceptance threshold schedules,
- web based visualizations of posterior parameter distributions, acceptance thresholds, configuration options and more,
- early stopping of model simulations, and
- a variety of configuration and custom extension options without alterations of pyABC’s source code.

pyABC can be combined with any user-defined computational model, distance function and parameter prior. Models can be defined as functions mapping the model parameters onto simulated data. This ensures a high degree of flexibility and allows the internal usage of, e.g., the Systems Biology Markup Language (SBML) or GPUs. Custom- and scipy.stats-distributions are supported as priors. Post-processing and analysis is supported via the included API which provides pandas data frames, or by directly querying the underlying relational database.

### 2.2 Multi-core and distributed execution

Single-machine multi-core execution and multi-machine distributed execution in cloud and cluster environments is featured by pyABC. A variety of distributed execution engines is supported, such as ad hoc clusters (e.g., the Dask distributed cluster and the IPython parallel cluster), bare grid-submission systems (e.g., SGE and UGE), and Redis based, low-latency setups. Furthermore, two parallelization strategies for the sampling of particles in the individual populations are provided:

- *Static Scheduling (STAT):* For each particle, one task is started on the available infrastructure. Within each task, proposal parameters are sampled and model simulations are run until exactly one simulation’s parameter is accepted (Fig. 1a,b). Denoting by n the number of desired particles, then even for infrastructures with more than n cores, only n cores are employed. The tasks are queued, if n is larger than the number of cores and are executed as slots become available. STAT aims to minimize the total amount of computation.
- *Dynamic Scheduling (DYN):* Parameter sampling and model simulation is continuously performed on all available hardware until n particles are accepted (Fig. 1c,d). All running simulations are then waited for to complete, yielding m ≥ n accepted particles. The n accepted particles started first are included in the next population while the remaining m–n accepted particles are discarded (to prevent bias towards parameters with shorter simulation times). DYN provides a scalable parallelization strategy aiming to minimize the total runtime.

For STAT, the degree of parallelism is limited to the population size n and decreases as particles are accepted (Fig. 1a), whereas DYN uses all available cores until n particles are accepted (Fig. 1c). Moreover, DYN scales further if the number of available cores is larger than the population size (Fig. 1e). When communication time is negligible compared to model simulation time, DYN is faster than STAT (Fig. 1e), however, STAT performs less computation overall (Fig. 1f).

**Figure 1:**
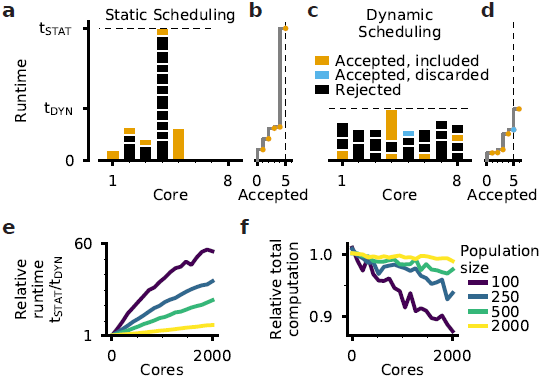
Parameter inference and scalability of pyABC’s parallelization strategies. (a-d) Static Scheduling (STAT) and Dynamic Scheduling (DYN) for 5 particles and 8 cores. The core usage (a, c) and the total number of accepted particles (b, d) are depicted over runtime. The color-coding indicates whether a sample satis1es the acceptance threshold and is included in the next population, satis1es the acceptance threshold but is discarded (i.e. not included in the next population) or does not satisfy the acceptance threshold and is rejected. In (c), the 1fth accepted sample (light-blue) is not included in the next population. (e) Mean relative runtime and (f) mean relative total amount of computation of STAT compared to DYN for exponentially distributed simulation times disregarding potential communication overhead.

### 2.3 Configuration, customization and extension

The pyABC package is modular and extensible facilitating to experiment with and to develop new ABC-SMC schemes. Following the documented API, transition kernels, (adaptive) acceptance threshold schedules, distance functions, summary statistics and other options can be customized and configured. Model simulation and distance calculation can be combined to interrupt the simulation and reject it early to reduce the runtime, e.g., if the distance is a cumulative sum as it is commonly the case for time series simulations. The framework can be run on new parallel environments providing corresponding custom map functions or implementations of the concurrent.futures.Executor interface. All customizations are possible without modifying pyABC’s source code.

## 3 Conclusion

pyABC addresses the need for distributed, likelihood-free inference for computationally demanding models. While pyABC’s less scalable STAT strategy is also implemented elsewhere (Jennings and Madigan 2017), the runtime optimized, more scalable DYN strategy is, to the authors’ knowledge, not available in any other Python ABC-SMC package. The flexibility and extensibility of pyABC render it applicable to a broad range of problem classes and infrastructures. We expect it to be used in many fields, including the emergent 1eld of multi-scale modeling.

